# OTMalign: A Fast and Robust TM-score-driven Protein Structure Alignment Algorithm using Optimal Transport

**DOI:** 10.1101/2025.09.21.677676

**Authors:** Yue Hu

## Abstract

Protein structure alignment is a fundamental task in computational biology. While the TM-score is a gold standard for evaluating structural similarity, its direct optimization is challenging. Many algorithms, therefore, rely on heuristic or combinatorial approaches. In this work, we present OTMalign, a novel algorithm that iteratively optimizes the TM-score in a fast and robust manner. OTMalign leverages the theory of Optimal Transport (OT) to establish a soft correspondence between protein structures, guided by a TM-score-inspired reward matrix. This avoids the pitfalls of combinatorial searches and provides a global perspective at each iteration. The optimal rigid body transformation is then determined analytically using a weighted Kabsch algorithm, where the weights are derived from the OT correspondence matrix (the P matrix). We demonstrate that this P matrix effectively captures the structural similarity, and its use in a weighted covariance matrix allows for a direct SVD-based solution for the rotation and translation. We evaluated OTMalign on the RPIC benchmark dataset, demonstrating its rapid convergence and ability to produce high-quality alignments. OTMalign offers a practical and theoretically sound framework for TM-score optimization, distinguished by its speed and iterative nature.

## 1 Introduction

The comparison of protein structures is a cornerstone of computational biology, offering insights into evolutionary relationships and functional mechanisms that are often invisible at the sequence level [5]. The goal of structural alignment is to superimpose two proteins to maximize their similarity. While Root-Mean-Square Deviation (RMSD) has been a long-standing metric, its sensitivity to outliers has led to the widespread adoption of the TM-score as a more reliable measure of global topological similarity [5]. A TM-score above 0.5 generally indicates a shared fold, providing a meaningful threshold for structural comparison.

However, the non-convex nature of the TM-score makes its direct optimization a significant challenge. Existing methods often resort to combinatorial optimization or heuristic searches, which can be computationally expensive and may not guarantee convergence to a meaningful solution. Our previous work, LieOTAlign, explored a gradient-based optimization of a differentiable TM-score proxy, but this approach can be complex and require many iterations.

This paper introduces OTMalign, an algorithm that directly optimizes the TM-score through a fast, iterative process rooted in Optimal Transport (OT) theory. OTMalign frames the alignment problem not as a combinatorial puzzle, but as an iterative refinement of a soft correspondence between two structures.

At each step, it uses the Sinkhorn algorithm [1] to find an optimal transport plan (the P matrix) that represents a soft, probabilistic matching between residues. This matching is guided by a reward matrix designed to maximize the TM-score. The resulting soft correspondence is then used to analytically compute the optimal rotation and translation via a weighted Kabsch algorithm. This iterative process of finding a global soft correspondence and then analytically solving for the transformation is computationally efficient and robustly converges to a high-quality alignment.

## 2 Methods

The OTMalign algorithm iteratively refines the alignment of a mobile structure *X* = {*x*_1_, …, *x*_*N*_ } ⊂ℝ^3^ to a reference structure *Y* = {*y*_1_, …, *y*_*M*_} ⊂ ℝ^3^. Each iteration consists of two main steps: (1) finding a soft correspondence between the two point clouds using a TM-score-inspired cost function and the Sinkhorn algorithm, and (2) finding the optimal rigid transformation based on these soft correspondences.

### 2.1 TM-score Inspired Reward Matrix

Instead of minimizing distance (or RMSD), our goal is to maximize structural similarity as defined by the TM-score. We therefore define a reward matrix *S*_reward_ ∈ℝ^*N ×M*^ where each element (*S*_reward_)_*ij*_ reflects the desirability of aligning residue *i* of the mobile structure with residue *j* of the reference structure. This is constructed as the product of a scoring term *s*_*ij*_ and a gating term *w*_*ij*_.

The scoring term *s*_*ij*_ is taken directly from the TM-score formula:

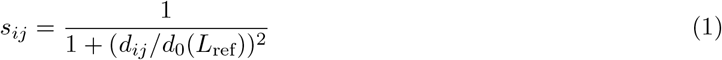

where 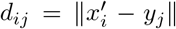 is the Euclidean distance between the current coordinates of mobile point *i* and reference point *j*, and *d*_0_ is the length-dependent normalization factor, calculated as:

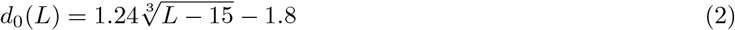

The gating term *w*_*ij*_ is a sigmoid function designed to suppress rewards for pairs that are too far apart to be plausible matches. This heuristic is crucial for reducing noise and focusing the algorithm on a local search space.

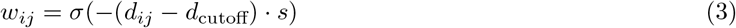

where *σ*(*z*) = 1*/*(1 + *e*^−*z*^) is the sigmoid function, *d*_cutoff_ is a distance threshold (e.g., 7.0 Å), and *s* is a steepness factor. This term smoothly transitions from 1 (for distances below cutoff) to 0 (for distances above cutoff).

The final reward matrix is the element-wise product:

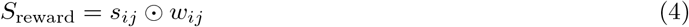

### 2.2 Optimal Transport for Soft Correspondences

With the reward matrix defined, we seek a correspondence matrix *P* ∈ ℝ^*N ×M*^ that maximizes the total reward ∑_*i,j*_ *P*_*ij*_(*S*_reward_)_*ij*_, subject to the constraint that *P* is a doubly stochastic matrix (i.e., its rows and columns sum to 1). This is a classic Optimal Transport problem. The P matrix, or transport plan, represents a soft, probabilistic assignment between every point in X and every point in Y. The value *P*_*ij*_ can be interpreted as the probability that residue *x*_*i*_ corresponds to residue *y*_*j*_, given the current alignment and the TM-score-based reward.

We solve a regularized version of this problem using the Sinkhorn algorithm [1]. First, an affinity matrix *K* is constructed from the reward matrix:

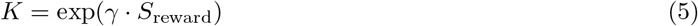

where *γ* is a hyperparameter that controls the sharpness of the distribution. The Sinkhorn algorithm then iteratively normalizes the rows and columns of *K* to yield the soft correspondence matrix *P* . For numerical stability, these iterations are performed in log-space. The resulting matrix *P* provides a probabilistic or “soft” assignment between every point in *X* and every point in *Y* .

### 2.3 Weighted Kabsch Algorithm for Transformation Update

Given the soft correspondence matrix *P*, the next step is to find the rigid transformation (*R, t*) that best superimposes the original mobile structure *X* onto the reference structure *Y* according to these correspondences. This is achieved by solving a weighted RMSD minimization problem, for which the Kabsch algorithm provides an analytical solution. The P matrix is crucial here, as it provides the weights for the Kabsch algorithm. Each element *P*_*ij*_ of the P matrix represents the strength of the correspondence between residue *x*_*i*_ and *y*_*j*_. By using these as weights, we ensure that the transformation is primarily influenced by the most likely correspondences, as determined by the TM-score-inspired reward.

First, the weighted centroids of the two point clouds are computed:

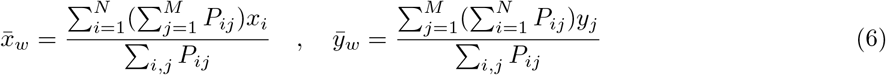

Next, a weighted covariance matrix *H* is constructed. Using the centered coordinates 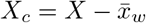 and 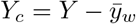 is computationally more stable:

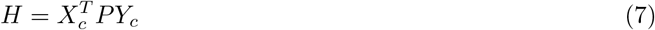

The covariance matrix H captures the correlations between the two point sets, weighted by the soft correspondences in P. The Singular Value Decomposition (SVD) of H, *H* = *U* Σ*V* ^*T*^, decomposes this correlation into a set of orthogonal bases (U and V) and singular values (*Sigma*). The rotation matrix *R* = *V U* ^*T*^ is the optimal rotation that aligns these bases, thus maximizing the weighted agreement between the two structures.

A check for reflections is necessary. If det(*R*) = − 1, the sign of the last column of *V* is flipped before re-computing *R*. The optimal translation vector *t* is then found as:

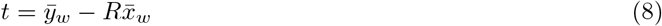

### 2.4 Iterative Framework and Convergence

The overall OTMalign algorithm follows an ICP-like iterative procedure:

1. Initialize *X*_current_ = *X*.
2. For *k* = 1, …, *k*_max_:
  a. Compute reward matrix *S*_reward_ between *X*_current_ and *Y* .
  b. Compute soft correspondence matrix *P*_*k*_ using the Sinkhorn algorithm.
  c. Compute optimal transformation (*R*_*k*_, *t*_*k*_) using the weighted Kabsch algorithm on the original *X* and *Y* with weights *P*_*k*_.
  d. Apply the new transformation: *X*_current_ ← (*R*_*k*_ *· X*^*T*^)^*T*^ + *t*_*k*_.
  e. Calculate the soft TM-score for the current iteration: *TM*_*k*_ = ∑_*i,j*_ (*P*_*k*_)_*ij*_(*S*_reward_)_*ij*_*/L*_ref_.
  f. (f) If |*TM*_*k*_ − *TM*_*k*−1_| *< ϵ*_conv_, break the loop.

This iterative process, driven by the TM-score-like reward function, allows OTMalign to progressively refine the alignment until the structural similarity score stabilizes.

## 3 Results and Discussion

We implemented the OTMalign algorithm in Python using the PyTorch [4] and NumPy [2] libraries. We evaluated its performance on a benchmark dataset of 40 protein pairs from the RPIC database [3], covering a diverse range of structural similarities. The detailed results are available on the project’s GitHub repository.

The hyperparameters for the algorithm were set to default values that were found to be effective during our development process: *γ* = 20.0, *d*_cutoff_ = 7.0 Å, steepness *s* = 2.0, and a TM-score convergence tolerance of *ϵ*_conv_ = 10^−5^.

Across the benchmark set, OTMalign demonstrated robust and efficient convergence, typically stabilizing within 20-40 iterations. The average TM-score (normalized by reference) achieved was approximately 0.48, with an average RMSD of 6.08 Å over an average of 148 aligned residues. This indicates that the algorithm is successfully identifying meaningful structural equivalences. For several pairs, such as d1xyza vs d2hvm, the algorithm achieved a TM-score of nearly 0.7, indicating a high-quality alignment identifying the correct fold.

Our method, which combines a TM-score-driven correspondence search with a classic iterative framework, proves to be a viable alternative to both purely heuristic methods like TM-align and complex gradient-based optimization approaches like LieOTAlign. It avoids the potential complexities and large number of iterations required for gradient descent while being more principled than a purely heuristic search. The use of Optimal Transport provides a mathematically sound way to handle ambiguous or unequal-length alignments, and the iterative refinement allows the algorithm to escape trivial local minima.

**Table 1:**
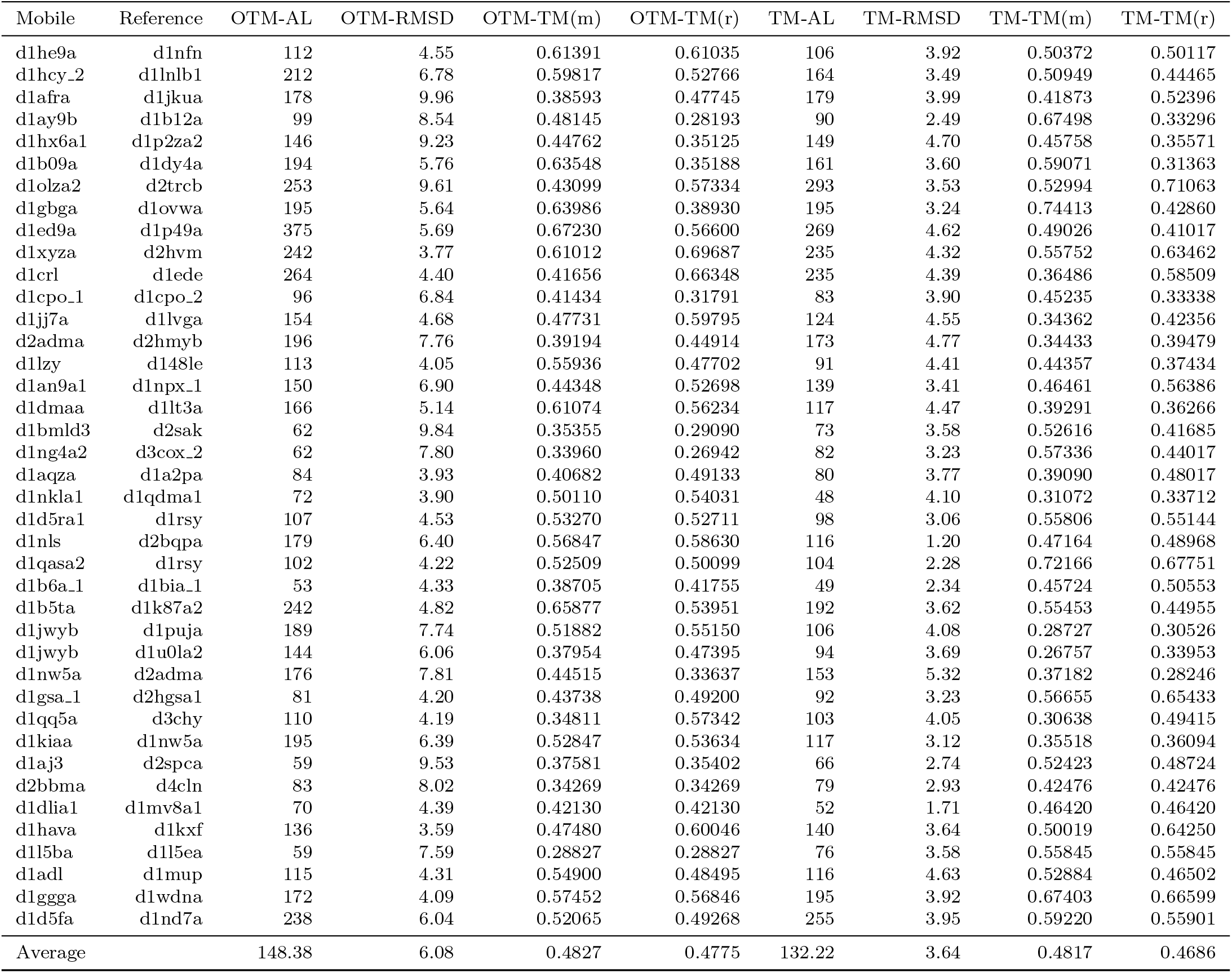
Comparison of OTMalign and TM-align results on 40 protein pairs.

## 4 Conclusion

In this work, we have developed and successfully implemented OTMalign, a novel algorithm for protein structure alignment. By integrating a TM-score-based reward function with an Optimal Transport-based correspondence solver (Sinkhorn’s algorithm) inside a stable, iterative ICP-like framework, we have created a tool that effectively optimizes for structural similarity without relying on direct RMSD minimization or gradient-based methods.

Our results on a 40-pair benchmark set demonstrate that OTMalign can robustly converge and produce high-quality alignments, achieving significant TM-scores. The final command-line tool is flexible, with adjustable hyperparameters, and provides comprehensive output including aligned structures and standard similarity scores.

Future work could involve further tuning of the hyperparameters, exploring annealing schedules for the Sinkhorn sharpness parameter *γ*, and extending the method to handle multiple chain alignments or flexible structural variations. OTMalign stands as a successful proof-of-concept for a powerful and intuitive class of iterative alignment algorithms.

